## Supplementary for "Epigenetic age acceleration in frontotemporal lobar degeneration: a comprehensive analysis in the blood and brain"

**Supplementary table S1.** Cohort description and characteristics.

| A) **Peripheral Blood** | | | | | | | | | | | | | | | | | | | | |
| --- | --- | --- | --- | --- | --- | --- | --- | --- | --- | --- | --- | --- | --- | --- | --- | --- | --- | --- | --- | --- |
| **Sample Group** | **No. of individuals** | | **Average Chronological age (SD)** | | **Average DNAm age (SD)** | | | | | | **Average age acceleration (residuals)** | | | | | | | | | |
| **Cohort 1** |  | |  | | **DNAmClock_Multi_** | **DNAmClock_Hannum_** | | **PhenoAge** | | **GrimAge** | **DNAmClock_Multi_** | | **DNAmClock_Hannum_** | | **PhenoAge** | | **GrimAge** | **IEAA_Multi_** | **IEAA_Hannum_** | **EEAA** |
| Controls | 178 | | 68.9 (10.4) | | 59.8 (10.5) | 68.1 (10.1) | | 58.4 (10.6) | | 59.8 (10.5) | -0.5 | | -1.2 | | -0.9 | | -0.6 | -0.5 | -0.8 | -1.6 |
| FTD | 117 | | 65.2 (9.0) | | 57.1 (9.3) | 66.7 (8.5) | | 57.4 (10.8) | | 57.1 (9.3) | 0.3 | | 0.8 | | 1.1 | | 0.6 | 0.4 | 0.7 | 1.1 |
| PSP | 44 | | 69.9 (7.3) | | 63.9 (9.4) | 73.9 (10.5) | | 59.6 (10.0) | | 63.9 (9.4) | 1.6 | | 3.0 | | -0.1 | | 0.8 | 1.0 | 1.7 | 3.7 |
| Total | 339 | | 67.7 (9.8) | | 59.4 (10.1) | 68.3 (9.9) | | 58.2 (10.6) | | 59.4 (10.1) |  | |  | |  | |  |  |  |  |
| B) **Post-mortem brain tissue** | | | | | | | | | | | | | | | |  |  |  |  |  |
| **Sample Group** | | **No. of individuals** | | **Average Chronological age (SD)** | | | **Average DNAm age (SD)** | | | | | **Average age acceleration (residuals)** | | | |  |  |  |  |  |
| **Cohort 2** | |  | |  | | | **DNAmClock_Multi_** | | **DNAmClock_Cortical_** | | | **DNAmClock_Multi_** | | **DNAmClock_Cortical_** | |  |  |  |  |  |
| Controls | | 8 | | 75.8 (5.6) | | | 76.1 (4.1) | | 91.2 (6.6) | | | 1.3 | | 0.0 | |  |  |  |  |  |
| FTLD-TDPA | | 7 | | 66.9 (4.8) | | | 65.8 (4.0) | | 81.4 (5.8) | | | -2.2 | | -0.6 | |  |  |  |  |  |
| FTLD-TDPC | | 8 | | 72.9 (4.8) | | | 73.2 (5.8) | | 88.7 (6.4) | | | 0.6 | | 0.5 | |  |  |  |  |  |
| Total | | 23 | | 72.0 (6.1) | | | 72.0 (6.3) | | 87.3 (7.3) | | |  | |  | |  |  |  |  |  |
| **Cohort 3** | |  | |  | | |  | |  | | |  | |  | |  |  |  |  |  |
| Controls | | 14 | | 78.4 (11.8) | | | 71.3 (11.2) | | 83.2 (12.6) | | | -0.2 | | -0.2 | |  |  |  |  |  |
| FTLD-TDPA (*GRN)* | | 7 | | 64.6 (7.6) | | | 62.4 (7.2) | | 70.3 (5.9) | | | 2.5 | | 1.1 | |  |  |  |  |  |
| FTLD-TDPB (*C9orf72)* | | 13 | | 63.8 (8.2) | | | 59.4 (8.3) | | 67.8 (9.3) | | | 0.1 | | -0.5 | |  |  |  |  |  |
| FTLD-17 (*MAPT)* | | 13 | | 60.9 (7.6) | | | 56.0 (5.5) | | 65.5 (8.9) | | | -0.9 | | 0.1 | |  |  |  |  |  |
| Total | | 47 | | 67.5 (11.5) | | | 62.4 (10.3) | | 72.1 (12.2) | | |  | |  | |  |  |  |  |  |
| **Cohort 4** | |  | |  | | |  | |  | | |  | |  | |  |  |  |  |  |
| Controls | | 71 | | 76.0 (8.0) | | | 70.7 (6.6) | | 82.3 (735) | | | -0.5 | | -0.5 | |  |  |  |  |  |
| PSP | | 93 | | 71.6 (5.3) | | | 68.6 (6.3) | | 79.7 (6.1) | | | 0.4 | | 0.4 | |  |  |  |  |  |
| Total | | 164 | | 73.5 (6.9) | | | 69.5 (6.5) | | 80.8 (6.9) | | |  | |  | |  |  |  |  |  |

FTD – frontotemporal dementia; PSP – progressive supranuclear palsy; FTLD-TDPA/C – FTLD with 43 kDa transactive response DNA-binding protein (TDP-43) positive inclusions, types A and C; *C9orf7­2* – *C9orf7­2* mutation carriers; *GRN* – *GRN* mutation carriers, FTLD-Tau – FTLD with tau-positive inclusions; *MAPT* – *MAPT* mutation carriers, SD – standard deviation, IEAA – intrinsic epigenetic age acceleration; EEAA – extrinsic epigenetic age acceleration.

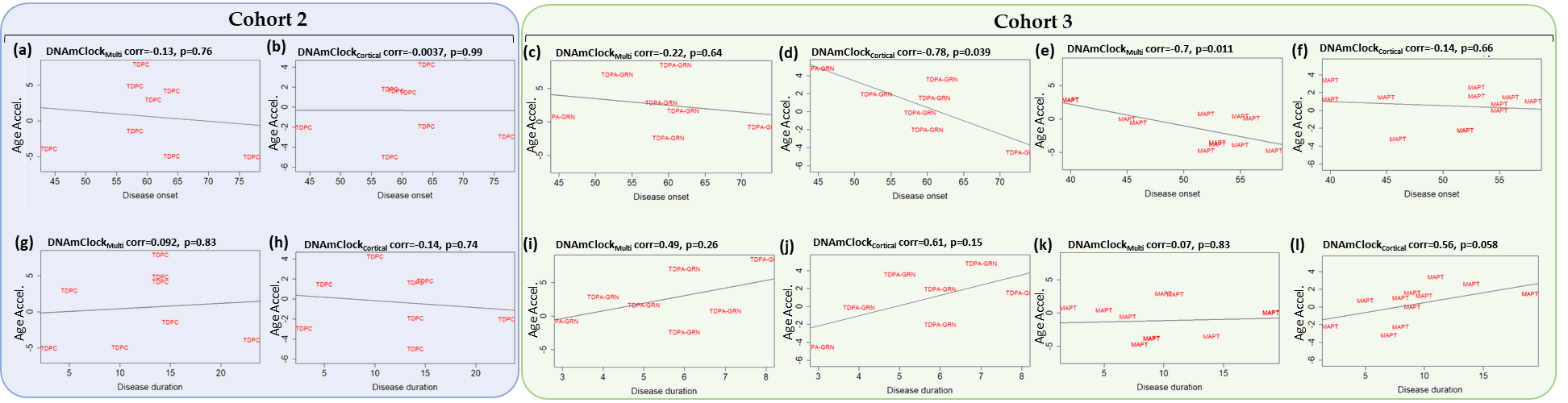

**Supplementary Figure S1.** Association between age acceleration and disease onset and duration with DNAmClock_Multi_ and DNAmClock_Cortical_ for the FTLD-TDPC subtype in cohort 2 and FTLD-TDPA-*GRN* and FTLD-Tau-*MAPT* mutation carriers in cohort 3: Age acceleration residuals (y-axis) for DNAmClock_Multi_ and DNAmClock_Cortical_ versus disease onset (x-axis) for (a,b) Cohort 2 (FTLD-TDPC), and (c-f) Cohort 3 (*GRN* and *MAPT* mutation carriers). Age acceleration residuals (y-axis) for DNAmClock_Multi_ and DNAmClock_Cortical_ versus disease duration (x-axis) for (g,h) Cohort 2 (FTLD-TDPC), and (i-l) Cohort 3 (*GRN* and *MAPT* mutation carriers). Age acceleration residuals were obtained by regressing DNA methylation age against chronological age and adjusting for confounding factors such as neuronal proportions obtained using a DNA methylation-based cell-type deconvolution algorithm. The correlation coefficient and p-values shown were calculated using Pearson correlation. TDPC – FTLD-TDP subtype C, TDPA-GRN – FLTD-TDPA *GRN* mutation carriers, MAPT – FTLD-Tau *MAPT* mutation carriers.
